# Patient-dependent epilepsy seizure detection using random forest classification over one-dimension transformed EEG data

**DOI:** 10.1101/070300

**Authors:** Marco A. Pinto-Orellana, Fábio R. Cerqueira

## Abstract

This work presents a computational method for improving seizure detection for epilepsy diagnosis. Epilepsy is the second most common neurological disease impacting between 40 and 50 million of patients in the world and its proper diagnosis using electroencephalographic signals implies a long and expensive process which involves medical specialists. The proposed system is a patient-dependent offline system which performs an automatic detection of seizures in brainwaves applying a random forest classifier. Features are extracted using one-dimension reduced information from a spectro-temporal transformation of the biosignals which pass through an envelope detector. The performance of this method reached 97.12% of specificity, 99.29% of sensitivity, and a 0.77 *h*^−1^ false positive rate. Thus, the method hereby proposed has great potential for diagnosis support in clinical environments.

## 1. Introduction

Epilepsy is the second most common neurological disease in humans after stroke (Guo et al., 2010). It is estimated between 40 and 50 million of patients suffering from this condition worldwide, which means 1% of the total world population (Tzallas et al., 2009; Guo et al., 2010; Fatichah et al., 2014). Although there are several medical treatments, 30% of the patient population have not a positive response to medication (Orosco et al., 2016) requiring expensive and long diagnosis processes (Das et al., 2016).

Seizures are the typical indicators for epilepsy diagnosis (Tzallas et al., 2009). A seizure is an abnormal excessive and hyper-synchronized neural activity in the brain (Tzallas et al., 2009; Das et al., 2016; Orosco et al., 2016) and it can be seen through electrical variations recorded from the whole brain mass or specific sections on its structure (Sierra-Marcos et al., 2015).

Among several imaging techniques of biosignals with medical relevance (Teplan, 2002), electroencephalograms (EEGs) are some of the most relevant alternatives. EEGs are recordings of electrical time-dependent variations of the brain activity (Teplan, 2002; Djemili et al., 2016). Although the electrical amplitude measurable in each neuron is small, the result of the synchronization in time and phase of large neural networks during cognitive operations allows detecting the neuron electrical signals even at scalp with EEG (Hesse et al., 2003).

There are several methods to recognize seizures using electrical biosignals of the brain (Tzallas et al., 2009). It was proved that epilepsy seizures are more distinguishable with electrical recordings in brain internal layers or electrocorticograms (ECoGs). Hence, EEG contains several distortions of the epilepsy signatures in comparison with ECoG (Spyrou et al., 2016). Nevertheless, there are several advantages of the former over other techniques: It is safe, non-invasive, and easy to assembly (Hu et al., 2011). Those characteristics established EEG as the *defacto* standard, along with video monitoring for epilepsy seizure diagnosis (Page et al., 2015).

Medical recommendations for seizure diagnosis often include performing long brain activity recordings of patients (Tsiouris et al., 2015). Then, the obtained data are analyzed by an expert in the area relying on subjective inspection (Samiee et al., 2015; Tsiouris et al., 2015; Spyrou et al., 2016; Orosco et al., 2016) commonly based on the visual information available (Das et al., 2016; Orosco et al., 2016). This human dependency implies an expensive and time-consuming process prone to errors due to the stored data size (Tsiouris et al., 2015).

It is important to notice that not all epilepticform waves appear during seizure intervals (Sierra-Marcos et al., 2015). Paroxysmal activity is an abnormal synchronous discharge of large ensemble of neurons and is strongly related with seizure processes. Such activity can be detected in EEG and is usually confused as an effectively epilepsy marker (Tzallas et al., 2009). This abnormal cortical pattern can appear during seizures (ictal) or in the interval between them (interictal) (Tzallas et al., 2009; Syrou et al., 2016). One of the bases of the present study is to explore and use that uncommon intorictal patterns as seizure predictor factors.

Several alternatives have been developed to recognize seizures in EEG (Orosco et al., 2013; Alotaiby et al., 2014). A first distinction between the approaches is the specific domain of the EEG data on which they have focused: Time series, frequency data, or spectro-temporal signals (Ahmad et al., 2015; Das et al., 2015). Another significant difference is the set of features which each method obtained in its analysis: Some studies utilized statistical, chaos theory. or information theory parameters (Ahmad et al., 2015; Das et al., 2015; Gill et al., 2015). while other efforts applied data transformations such as singular value decomposition (SVD) or principal components analysis (PCA) (Zhao et al., 2016). A third prominent divergence among the previously proposed methods is the level of artificial intelligence dependency to perform a decision. A group of methods was designed to detect epilepsy intervals without needing classifiers based on machine learning (Alotaiby et al., 2014), while other studies required specific algorithms to classify well the data. In the latter category, support vector machines (SVM), k-nearest neighbours (KNN), and artificial neural networks (ANN) are frequently used as classifiers (Orosco et al., 2013; Alotaiby et al., 2014).

This study proposes a novel approach which obtained a higher performance compared with other state of the art studies. This method is a combination of several techniques of signal processing and machine learning: Short time Fourier transform, principal component analysis, maximum moving filter and a random forest classifier.

## 2. Materials

In this research, the data source relies on the CHB-MIT electroencephalographic scalp database from the Physionet project (Goldberger et al., 2000). The database stores intractable epileptic seizures from several pediatric patients at the Children’s Hospital Boston (Shoeb et al., 2004; Shoeb and Guttag, 2010). The data were collected for an experiment which was conducted for monitoring patients after withdrawal of the epileptic medication as an analysis before a surgical intervention. The dataset was collected in 23 cases from 22 different subjects where one of them was recorded again after 1.5 years.

The complete data package is compounded of 686 files saved in the European data format (EDF) representing a total of 961.64 hours (Goldberger et al., 2000). The seizure recording durations of each patient are detailed in Table 1. All signals were measured with a sampling frequency of 256Hz with 16 bits of resolution. Nearly all files were captured in 23 channels using the international 10-20 system of electrode positions and nomenclature.

**Table 1:** Recorded seizure durations of each patient. It should be noted that chb02 and chb24 represent recordings of the same patient but with a difference of 1.5 years between the recording dates. However, they are identified as different subjects for the experiment.

| Patient ID | Number of seizures | Seizure average durations | Seizure maximum durations | Seizure minimum durations |
| --- | --- | --- | --- | --- |
| chb01 | 7 | 63.14 | 101 | 27 |
| chb02 | 3 | 57.33 | 82 | 9 |
| chb03 | 7 | 57.43 | 69 | 47 |
| chb04 | 4 | 94.5 | 116 | 49 |
| chb05 | 5 | 111.6 | 120 | 96 |
| chb06 | 10 | 15.3 | 20 | 12 |
| chb07 | 3 | 108.33 | 143 | 86 |
| chb08 | 5 | 183.8 | 264 | 134 |
| chb09 | 4 | 69 | 79 | 62 |
| chb10 | 7 | 63.86 | 89 | 35 |
| chb11 | 3 | 268.67 | 752 | 22 |
| chb12 | 40 | 36.88 | 97 | 13 |
| chb13 | 12 | 44.58 | 70 | 17 |
| chb14 | 8 | 21.12 | 41 | 14 |
| chb15 | 20 | 99.6 | 205 | 31 |
| chb16 | 10 | 8.4 | 14 | 6 |
| chb17 | 3 | 97.67 | 115 | 88 |
| chb18 | 6 | 52.83 | 68 | 30 |
| chb19 | 3 | 78.67 | 81 | 77 |
| chb20 | 8 | 36.75 | 49 | 29 |
| chb21 | 4 | 49.75 | 81 | 12 |
| chb22 | 3 | 68 | 74 | 58 |
| chb23 | 7 | 60.57 | 113 | 20 |
| chb24* | 16 | 31.94 | 70 | 16 |

## 3. Method

This study proposes to use a mapping of a spectro-temporal transform of the brain signals into a one-dimension space for being used as input for a classifier algorithm (Figure 1). Thus, the complete signal analysis could be explained using four processes: Data preprocessing, dimensionality reduction, envelope detection, data regrouping, and classification.

**Figure 1:**
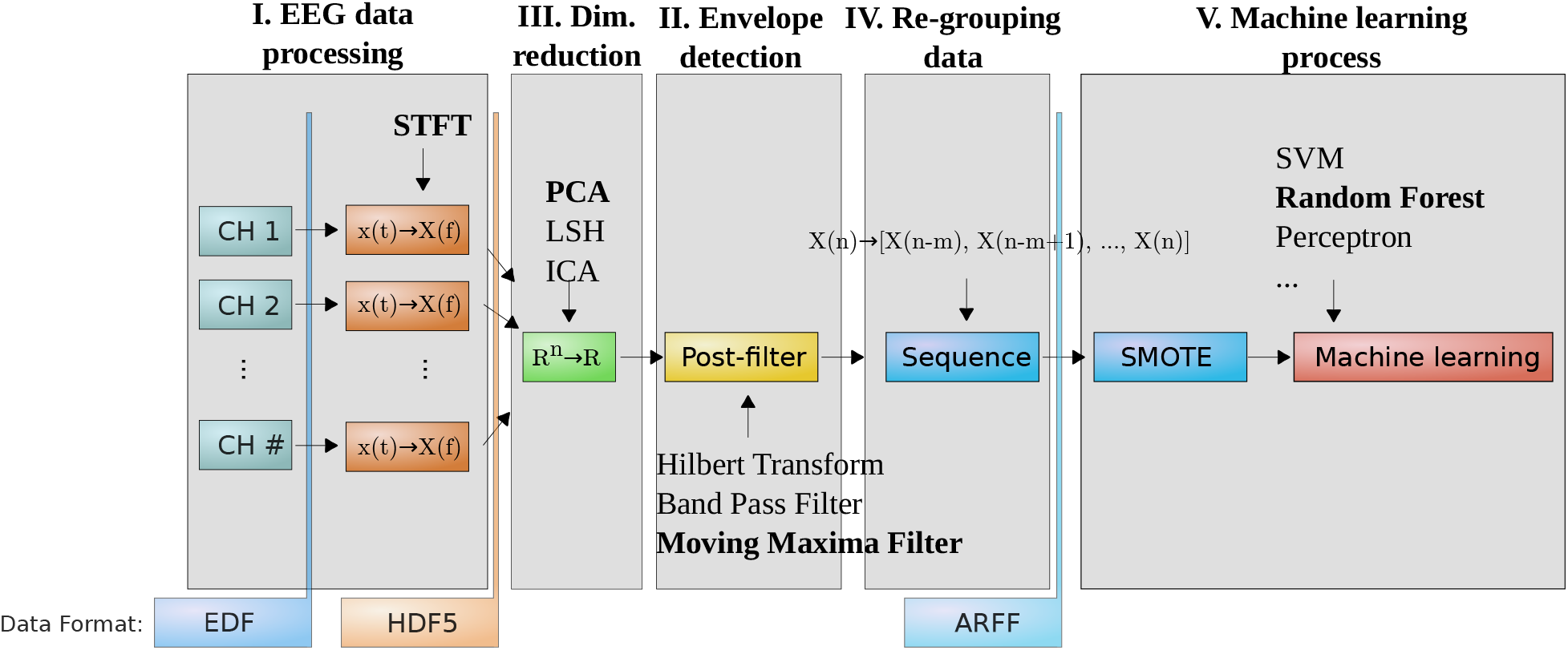
Workflow of the seizure detection process: Data preprocessing, dimensionality reduction, envelope detection, data regrouping, and classification.

### 3.1. EEG data processing

Initially, some characteristics must be explained about EEG. It exhibits several unique features: It is nonstationary, nonlinear, and frequency-variant. The nonstationary property denotes there is no period of time where the exact signal will be repeated (Natarajan et al., 2004; Yan et al.,2015). Also, as a non-linear time series (Natarajan et al., 2004), EEG does not allow superposition or homogenization, preventing anticipating future values using a linear combination of the previous ones:

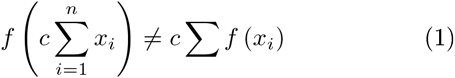

where *c* is a real constant,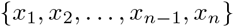 are the sampling instants, and *f* (*x*_*i*_) is the point related with a instant *x*_*i*_.

Associated to the last attribute, variations in the information source on different conditions or activities are linked with alterations in the spectral response of the brain biosignals (Yan et al., 2015).

Several techniques have been developed to extract information of EEG time series. The majority of them can be grouped in two categories: Nonlinear feature extraction processes and time-frequency transformations.

The first group is comprised of the nonlinear analysis techniques based on information and chaos theory tools. This kind of methods use parameters which can be used successfully with non-linear and multivariate time series (Natarajan et al., 2004). The most frequent feature in this category is the signal entropy, which is a thermodynamic-inspired quantity to describe the amount of information (Awan et al., 2015; Iqbal et al., 2015). The Hurst exponent, which measures the signal self-similarity, allowing detecting time dependences, is often used as another statistical feature. Another parameter, the Lyapunov exponent, which measures the sensitive dependence of the data regarding to their initial conditions, is a representative of the chaos theory derived metrics (Natarajan et al., 2004).

The second group is made of spectro-temporal transformations. Information in the brain electrical signals is stored with frequency variations (Yan et al., 2015). For this reason, it is useful to analyze the deviations which appear in the frequency domain. The conventional alternative is a discrete Fourier transform or a fast Fourier transform (Alotaiby et al., 2014). However, these kind of mathematical tools are focused in stationary data series and cannot be directly applied in EEG (Natarajan et al., 2004).

A typical approach to analyze nonstationary signals is splitting them in several time intervals, or time windows, such that it could be obtained spectral features while partial time information is retained (Tzallas et al., 2009). In this perspective, a direct method is short time Fourier transform (STFT), also known as Gabor transform, which splits the complete signal in several fixed-longt.il windows and applies a Fourier transform in each slot. A significant disadvantage of this method is the fixed time-froquency resolution. However, it showed an optimal response to detect seizures (Samiee et al., 2015; Yan et al., 2015). Alternative methods such as Wavelet transforms (WT) or Wigner-Ville distributions (WVD) allow to obtain better resolution. Nonetheless, either there is no a relationship between the local frequencies and the measured time (as in the case of WT) or nonlinear signals added to the transformed data (as in the case of WVD) (Yan et al., 2015).

Applying only frequency transforms over EEG data could lead to information loss due to their non-stationary condition (Al-Manie and Wang, 2015). Therefore, STFT was selected as the procedure to map signal into a time-froquency plane (Wang et al., 2015). For a given discrete function *x* [*n*], the STFT is calculated shifting a small sliding window *g* [*n*] of a Δt length over the time series to obtain the frequency spectrum in each time interval:

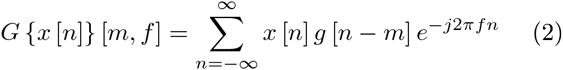

where *n* and *m* are sampling instants, and ƒ is the signal frequency.

Due to the fact that the signal is not continuous (it is sampled at fixed periods) there are several limitations in the process of STFT calculation. A first restriction is linked with its discrete nature, making the time resolution, which is the minimum time interval when frequency information is reteived,depend on the length of points Δ*n* and the window *g* [*n*]:

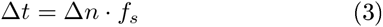

Another constraint is the frequency resolution. STFT only obtains frequency amplitudes with the form 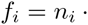 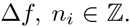 Additionally, the resolution in frequency Δƒ is calculated with the following expression:

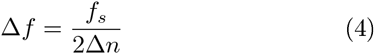

For the datasets described here, it was used a time window Δ*t* of 39 microseconds, which allowed a range of Δƒ 12.8 Hz.

### 3.2. Data, dimensionality reduction

Electroencephalograms are multivariate data and their computed features have a high dimensionality, 230 dimensions in this research, which difficults their visualization and processing (Birjandtalab et al., 2016). This problem, known as “the curse of dimensionality”, could impact the classifier performance increasing misinterpretations or over interpretation of the data (Ahmad et al., 2015). Common procedures to reduce these inconveniences rely on selection of the most significative features.

There are several methods previously used for reducing EEG data dimensionality: Principal component analysis (PCA) (Alotaiby et al., 2014), independent component analysis (ICA) (Vigário et al., 2000), T-distributed stochastic neighbor embedding (Birjandtalab et al., 2016), and local sensitive hashing (LSH) (Perronnin et al., 2010). After several experiments (not shown) of speediness and memory consumption, PCA was selected as the method of this study.

PCA is a technique which linearly transforms the original variables into other uncorrelated ones. These result variables, known as principal components, are orthogonal among them. The PCA transformation is performed in such a way that the first principal component accounts for the highest variablity in the data, the second principal component accounts for the second highest variablity in the data, and so on (Patel et al., 2015). Therefore, all components are assigned to a score which denotes the variance of the partial set they represent of the data (Czarnecki and Gustafsson, 2015).In this way, one can keep the components with highest scores to reduce data dimensionality.

In (Zhao et al., 2010), it was studied the influence of different mapping functions, or kernels, and PCA along with a support vector machine classifier with EEG data. It was shown three dimensions could reach up to 100% of accuracy in several conditions concerning the classifier settings, the type of mapping function, and the kind of mental state analyzed.

PCA was applied over the EEG multivariate time series maintaining only the component with maximum score. The aim of mapping to only one dimension is allowing to represent graphically the whole brain signal in one chart maintaining a relationship between the signal shape and the EEG epilepsy state.

### 3.3. Signal envelope detection

Previous experiments (not shown) have revealed several associations between the PCA reduced data and the seizure marks. This connection is apparently related to the PCA signal shape as it can be seen in Figure 2. It should be noted that this behavior corresponds to amplitude modulated signals. In consequence, it was applied an envelope detection process over the PCA signal. There are several methods available for this goal: Hilbert transform, pass band filters, and hysteresis transform. However, after a trade-off analysis between processing time and resultant shape, it was selected the simplest off-line method: The moving maxima filter.

**Figure 2:**
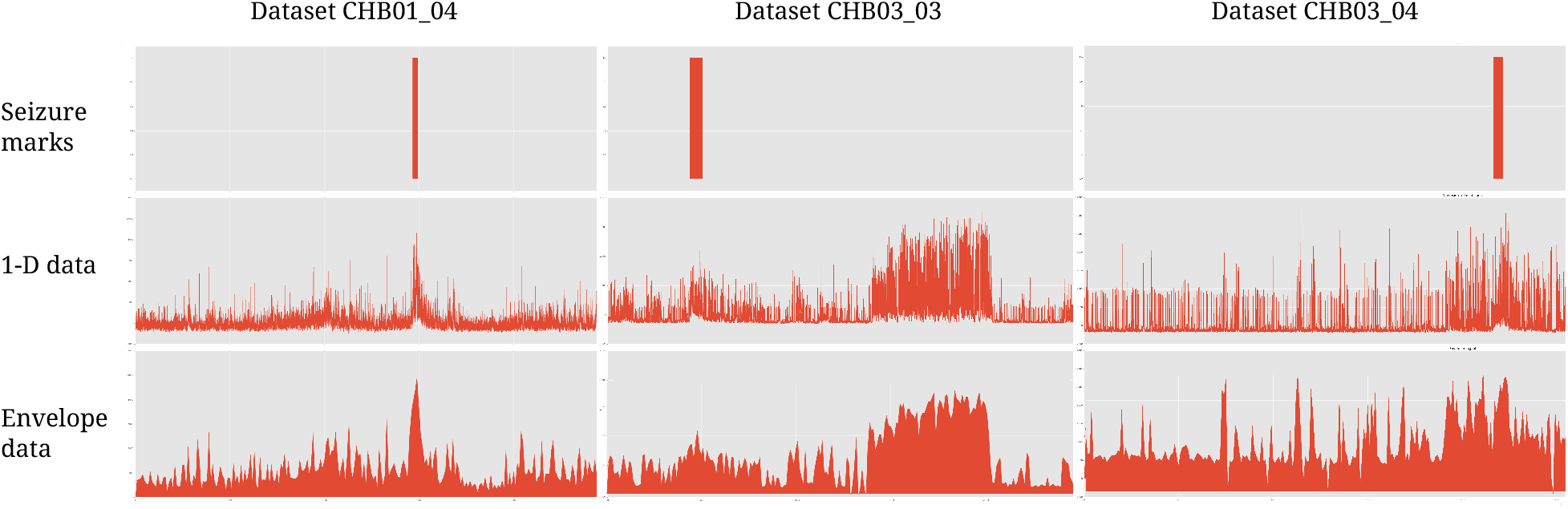
Graphical comparison between the PCA converted data and its envelope regarding the seizure marks in the datasetsCHB01_04,CHB03_03 and CHB03_04.

The moving maxima filter is a process that consists in creating a new series *h*[*n*] from an original series *g*[*i*] such that each point *n* the maximum value between *g*[*n*]and *g*[*n*+Δ*n*]:

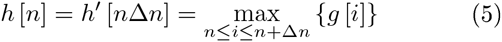

An equivalent method was developed in (Zhang and Smith, 2001). It had been applied to the analysis of evolution indexes in financial time series. As it will be explained later, the goal of this filter is finding the envelope of the signal in mechanism similar to amplitude demodulator, only on the time period Δ*n* which was defined as 10 seconds in the present project.

### 3.4 Re-sequencing frequency-time data

One of the assumed hypotheses of this study is considering the prediction of a current individual state as a variable dependent on the current data point and a fixed set of past values after the EEG signal processing:

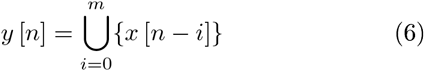

Following this argument, the enveloped data were reorganized in blocks of fixed size. The block size m could have an influence in the final system ability to detect a seizure state and it may be tested with several values. However, due to the limitations related with the data size, only two block sizes were selected: 30 and 70 seconds.

### 3.5. Machine learning: Balancing and classification

The last step in the method was applying a machine learning algorithm. Nevertheless, due to the typical imbalance of this kind of datasets (Tsiouris et al., 2015), the points marked as seizure comprise a very small set in comparison to the interictal intervals. In consequence, firstly, it was applied the SMOTE algorithm over the data for treating the imbalance problem. In (Fergus et al., 2015),this oversampling technique was used efectivelly in other EEG analysis to improve recognition over noise datapoints which were not considered in the original dataset.

After the application of the SMOTE method, a random forest classifier was configured and executed. Random forest is an ensemble learning algorithm which uses multiple decision trees (Shiratori et al., 2015). Its working method consists in creating a fixed number of random sets from the original training set using random feature selection and a bootstrapping method. For each new set, a different decision tree is trained without pruning. Thus, each instance decision is based on the majority vote obtained from the decision trees (Peker et al., 2015; Shiratori et al., 2015).

Several studies have shown that random forest works satisfactorily with featured extracted from EEG (Patti et al., 2015; Peker et al., 2015; Shiratori et al., 2015). Also, in the seizure-specific context, (Wang et al., 2015) found average accuracies greater than 96% using this classifier on the epilepsy dataset of the University of Börn.

In this study, the random forest algorithm was configured to work with 100 decision trees without prune restriction. It was utilized the random forest implementation of the WEKA API 1.7.0 (Hall et al., 2009).

## 4. Results

The method proposed in this work was applied to the physiological signals stored in the CHB-MIT database. Every channel was processed with the STFT transform, and the dimensionality of the multivariate set was reduced using PCA. Then, it passed through an envelope detector and, finally, the detected envelope was reorganized in blocks of fixed length to be processed by a random forest classifier.

As it was mentioned previously, it was selected two possibilities at the re-sequencing process: Blocks of 30 and 70 seconds. Different sizes showed the influence of a seizure interval, ictal or interictal, regarding to the past values of the transformed signal.

Each defined configuration was tested and executed with the proper software tools, with 10-fold cross validation as evaluation method. Regarding the performance indexes, it was used standard parameters: accuracy (ACC), specificity (SPE), sensitivity (SEN), and false positive rate (FPRe) (Ahmad et al.,2015; Gill et al., 2015; Iqbal et al., 2015; Tsiouris et al., 2015; Das et al., 2016; Orosco et al., 2016; Zhao et al., 2016):

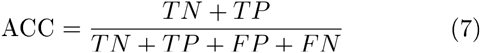

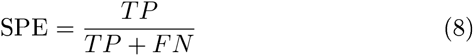

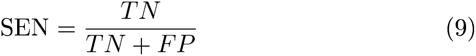

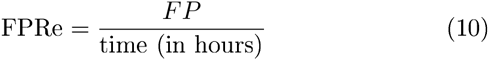

where TP is the number of true positives, TN is the quantity of true negatives, FP is the amount of false positives, FN is the number of false negatives, and time is a unit of analysis time. It should be noted that FPRe is measured in samples per hour (*h*^−1^).

Each data block, or instance, is considered positive (P) when it is within any of the seizure intervals that was labeled by the human specialists, or is identified as negative (N) when there is no seizure evidence in the EEG. Thus, the prediction is considered as a true negative or a true positive when the class defined by the algorithm matches. Hence, if the algorithm estimated a seizure event and the patient has no seizure, the prediction is flagged as a false positive, or in the other hand, as a false negative when the algorithm indicated an absent of seizure while there is evidence about the patient suffered it during that interval.

The obtained results were compiled in Table 2 and Table 3.The experiments showed a good performance with data blocks of 30 seconds, reaching average sensitivity of 89.73%, specificity of 94.77%, FPRe of 6.87 *h*^−1^, and accuracy of 92.46%. However, increasing the data block size to 70 seconds significantly improved the performance. The average values of the same measures were raised to: 97:12% of sensitivity, 99:29% of specifity, 98:30%of accuracy, and 0:77 *h*^−1^ of FPR.

**Table 2:** Performance evaluation parameters for each patient with a block size of 30 seconds

| Patient ID | ACC (%) | SEN (%) | SPE (%) | FPR/h |
| --- | --- | --- | --- | --- |
| chb01 | 96.55 | 94.38 | 98.38 | 2.27 |
| chb02 | 87.46 | 84.40 | 90.05 | 12.13 |
| chb03 | 96.50 | 93.74 | 98.84 | 1.53 |
| chb04 | 94.41 | 91.14 | 97.20 | 15.35 |
| chb05 | 96.25 | 95.61 | 96.79 | 4.33 |
| chb06 | 89.79 | 86.57 | 92.52 | 17.51 |
| chb07 | 88.56 | 85.95 | 90.78 | 21.68 |
| chb08 | 90.89 | 86.17 | 94.85 | 3.55 |
| chb09 | **98.41** | **98.38** | **98.44** | **3.71** |
| chb10 | 94.80 | 91.89 | 97.28 | 4.75 |
| chb11 | 96.39 | 94.21 | 98.24 | 2.12 |
| chb12 | **86.13** | **82.34** | **89.33** | **8.64** |
| chb13 | 92.62 | 90.15 | 94.71 | 6.03 |
| chb14 | 86.73 | 84.26 | 88.83 | 10.03 |
| chb15 | 89.13 | 83.62 | 93.78 | 8.57 |
| chb16 | 87.40 | 83.37 | 90.82 | 6.02 |
| chb17 | 93.84 | 89.55 | 97.47 | 1.75 |
| chb18 | 89.30 | 88.00 | 90.40 | 11.82 |
| chb19 | 96.08 | 97.10 | 95.22 | 4.95 |
| chb20 | 95.76 | 92.70 | 98.36 | 1.56 |
| chb21 | 95.62 | 92.29 | 98.46 | 1.75 |
| chb22 | 89.28 | 84.69 | 93.17 | 7.32 |
| chb23 | 94.28 | 91.19 | 96.90 | 2.88 |
| chb24 | 92.76 | 91.79 | 93.58 | 4.71 |
| **Average** | **92.46** | **89.73** | **94.77** | **6.87** |
| **Minimum** | 86.13 | 82.34 | 88.83 | 1.53 |
| **Maximum** | 98.41 | 98.38 | 98.84 | 21.68 |

**Table 3:** Performance evaluation parameters for each patient with a block size of 70 seconds

| Patient ID | ACC (%) | SEN (%) | SPE (%) | FPR/h |
| --- | --- | --- | --- | --- |
| chb01 | 99.62 | 99.24 | 99.95 | 0.16 |
| chb02 | 97.26 | 96.38 | 98.00 | 2.42 |
| chb03 | 98.76 | 97.38 | 99.94 | 0.08 |
| chb04 | 99.54 | 99.06 | 99.96 | 0.23 |
| chb05 | 99.82 | 99.73 | 99.91 | 0.13 |
| chb06 | 99.54 | 99.07 | 99.93 | 0.16 |
| chb07 | 99.27 | 98.69 | 99.77 | 0.53 |
| chb08 | 96.41 | 93.50 | 98.85 | 0.78 |
| chb09 | **99.90** | **99.85** | **99.94** | **0.14** |
| chb10 | 98.38 | 97.25 | 99.34 | 1.14 |
| chb11 | 98.81 | 97.83 | 99.63 | 0.44 |
| chb12 | 95.14 | 93.39 | 96.62 | 2.72 |
| chb13 | 97.30 | 95.83 | 98.54 | 1.65 |
| chb14 | 98.08 | 97.16 | 98.86 | 1.02 |
| chb15 | **95.00** | **91.67** | **97.80** | **3.00** |
| chb16 | 97.66 | 95.74 | 99.29 | 0.46 |
| chb17 | 97.77 | 95.34 | 99.83 | 0.12 |
| chb18 | 97.42 | 96.39 | 98.29 | 2.08 |
| chb19 | 99.85 | 99.80 | 99.89 | 0.12 |
| chb20 | 99.21 | 98.40 | 99.90 | 0.09 |
| chb21 | 99.11 | 98.16 | 99.93 | 0.08 |
| chb22 | 98.40 | 96.69 | 99.85 | 0.16 |
| chb23 | 98.25 | 96.43 | 99.78 | 0.20 |
| chb24 | 98.61 | 97.88 | 99.23 | 0.56 |
| **Average** | **98.30** | **97.12** | **99.29** | **0.77** |
| **Minimum** | 95.00 | 91.67 | 96.62 | 0.08 |
| **Maximum** | 99.90 | 99.85 | 99.96 | 3.00 |

## 5. Discussion

It should be noted that the database used in this paper is publicly available (Goldberger et al., 2000). As a result, there are other seizure detection methods that process the same information to evaluate their algorithms. It was selected 9 methods from 7 studies which analyzed the same databaseto compare with out approach. Their performances are compiled in Table 4, although some indexes were not found in their evaluation analysis. Among the alternative studies, several researches emphasized the importance of the sensibility in the evaluation over other values and only presented that value(Iqbal et al., 2015; Tsiouris et al., 2015).

**Table 4:**
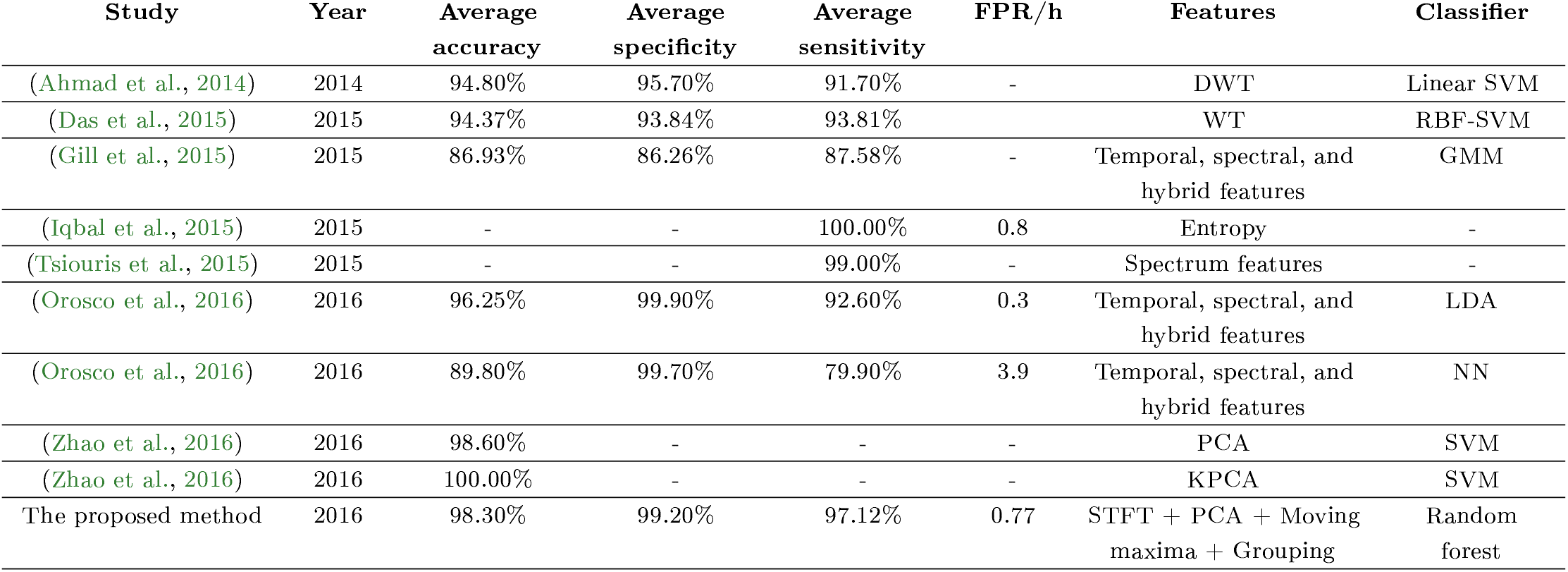
Performance comparison with other methods

| Study | Year | Average accuracy | Average specificity | Average sensitivity | FPR/h | Features | Classifier |
| --- | --- | --- | --- | --- | --- | --- | --- |
| (Ahmad et al., 2014) | 2014 | 94.80% | 95.70% | 91.70% | - | DWT | Linear SVM |
| (Das et al., 2015) | 2015 | 94.37% | 93.84% | 93.81% | - | WT | RBF-SVM |
| (Gill et al., 2015) | 2015 | 86.93% | 86.26% | 87.58% | - | Temporal, spectral, and hybrid features | GMM |
| (Iqbal et al., 2015) | 2015 | - | - | 100.00% | 0.8 | Entropy | - |
| (Tsiouris et al., 2015) | 2015 | - | - | 99.00% | - | Spectrum features | - |
| (Orosco et al., 2016) | 2016 | 96.25% | 99.90% | 92.60% | 0.3 | Temporal, spectral, and hybrid features | LDA |
| (Orosco et al., 2016) | 2016 | 89.80% | 99.70% | 79.90% | 3.9 | Temporal, spectral, and hybrid features | NN |
| (Zhao et al., 2016) | 2016 | 98.60% | - | - | - | PCA | SVM |
| (Zhao et al., 2016) | 2016 | 100.00% | - | - | - | KPCA | SVM |
| The proposed method | 2016 | 98.30% | 99.20% | 97.12% | 0.77 | STFT + PCA + Moving maxima + Grouping | Random forest |

The current research proposes a different method to process electroencephalographic signals with a high detection accuracy. Comparing with alternative methods,our technique showed the best performance. Sensitivity was just behind the technique based in an entropy-related parameter classification explained in (Iqbal et al., 2015)which obtained a 100% for SEN but did not present other comparable evaluation parameter.

Our approach obtained a slightly lower values concerning specificity and false positive rate (a difference of0:70% and 0:44 *h*^−1^) compared with one of the methods presented in (Orosco et al., 2016). However, in general the proposed technique demonstrated to be the best approach when all parameter values are taken into account in combination.

A good system response depends on the combination of the internal selected algorithms. Concerning feature selection, the majority of alternatives relies, at least partially,on spectral features: STFT, wavelet transform, or discrete Fourier transform (Ahmad et al., 2015; Gill et al.,2015; Tsiouris et al., 2015; Das et al., 2016; Orosco et al.,2016). Also, the dimension reduction using PCA or an improved algorithm based on it proved to reach accuracies close to 100%(Zhao et al., 2016). The present research combined both processing types using STFT over biosignals and then PCA on the time-spectrum data. The notable difference with respect to other studies is the use of a singular element typical of amplitude demodulators: An envelope detector to improve match between waveforms and seizure marks. Regarding the classification method,several authors used SVM (Ahmad et al., 2015; Das et al.,2016; Zhao et al., 2016), Bayessian classifers (Gill et al.,2015), linear discriminators or neural networks (Orosco et al., 2016).

Working eith other types of datasets, (Czarnecki and Gustafsson, 2015) denoted that a random forest classifier can obtain a better performance than other algorithms,including SVM, with reasonable less training time. Therefore,this machine learning algorithm was selected for our study.

## 6. Conclusions

This study describes a patient-dependent system for detecting seizures with a great prediction confidence. The structure of the proposed approach comprises a sequence of several signal processing and machine learning algorithms. The processing step includes a PCA process after a STFT transformation with a subsequent envelope shape detection stage. After those procedures, the new process of the data sequence involves a data regrouping in fixed length blocks that are then given as input to a random forest
classifier.

The parameters of the proposed system were optimized to maximize the overall performance. Thus, after training and testing with the CHB-MIT database containing 23 subject cases, it was obtained average values of 98:30% for accuracy, 97:12% for sensitivity, 99:29% for specificity, and 0:77 *h*^−1^ as false positive rate. Comparing these indexes with state of art alternative systems, we can conclude that a hardware implementation of our method could lead to a considerable positive impact on epilepsy diagnosis through the automation of seizure detection.

## 7. Acknowledgement

This work was supported by grants from Fundação de Amparo à Pesquisa de Minas Gerais (FAPEMIG) and the PAEC agreement between the Organization of American States (OAS) and Universidade Federal de Viçosa (UFV).

